# Eco-ethology and trait distribution of two congeneric species – different strategies for the plague pest *Acanthoplus discoidalis* and the long-legged *A. longipes* (Orthoptera: Tettigoniidae)

**DOI:** 10.1101/751826

**Authors:** Erminia Conti, Giovanni Costa, Christian Mulder

**Affiliations:** Department of Biological, Geological and Environmental Sciences, Section of Animal Biology, University of Catania, 95124 Catania, Italy

**Keywords:** *Acanthoplus* (Tettigoniidae), behaviour, body size, interspecific variance, intraspecific variance, trait distribution

## Abstract

High trait variability among insects reflects a combination of intra- and inter-phenotype variations. Our aim was to assess if the trait distribution of body measurements can be more significantly influenced by sex (intraspecific variance) or by species (interspecific variance). To achieve this, we collected in Namibia tettigoniids belonging to two congeneric species of armoured ground crickets: *Acanthoplus discoidalis* (a significant pest in African croplands) and the long-legged *Acanthoplus longipes*. We measured in the field the total body length, the maximal pronotal width and length, and the femur and tibia lengths of the hind legs in 106 adults. We also derived the body mass from length and width values of the sampled specimens. No significant differences emerged in the two species by sex. A discriminant analysis clearly shows that at species level the locomotory traits as captured by tibia and femur lengths and the size traits as captured by body and pronotal lengths account for 99% of the total variance and clearly separate this pest from its congeneric species. In essence, it is not primarily the body size that differentiates the two species, but rather the pronotum and hind leg larger sizes of *A. longipes*. Different eco-ethological requirements, like the peculiarity of the calling song and the movements within the vegetation (and the consequently needed energy), independently force these functional traits.

## INTRODUCTION

Extensive trait variability of morphological structures (legs, wings, abdomen, etc.) characterizes insect populations and mirrors behavioural adaptations from feeding and dispersal up to reproduction and competition. Hence it is important to investigate the many functional links between behavioural modifications and morphological traits, both at interspecific and intraspecific level (Violle *et al*. 2012). Path analysis by *F*-statistics (Wright’s 1949, 1965) was a first attempt to elucidate the pattern and the extent of genetic variation within and among natural populations. A population’s niche was suggested to be determined by the combination of two phenomena in resource use (Roughgarden 1972), i.e. an intra-phenotype variation and an inter-phenotype variation (Violle *et al*. 2012).

Such results are applicable in ecology and are particularly interesting if seen in the dynamic framework of interspecific competition: an understanding of differences in fundamental resources use by animals with different traits allows us to increase the effectiveness of dynamic forecasting of their reactions to their environment. From the perspective of a size-scaled behaviour, it is true that “a knowledge of allometry might allow animal behaviorists a greater definition of the probable behavior a study animal might exhibit” (Peters 1983). For instance, shifts in an insect consumer’s size will be mediated by type and rugosity of the habitat (Teuscher *et al*. 2009; Thakur *et al*. 2020) and by competitive strategies employed by that species (Amarillo[Suárez *et al*. 2011; Kalinkat *et al*. 2015; Hirt *et al*. 2018).

For instance, resource availability exerts a bottom–up control on leaf consumers (Lavorel *et al*. 2013; Mulder *et al*. 2012, 2013). We need to examine whether, for example, good resource quality simply favours species with larger body size or favours species with more aggressive competitive behaviour, typical of pest taxa (Sterner & Elser 2002; Mulder & Elser 2009). For insect locomotion the allometry of morphology is highly relevant (Kaspari & Weiser 2007; Dial *et al*. 2008; Whitman 2008; Kalinkat *et al*. 2015; Hirt *et al*. 2017, 2018; Thakur *et al*. 2020; see also Tamburello *et al*. 2015 for vertebrates). To understand the actual interrelationships between taxonomy, behaviour and functional traits in one pest and its congeneric species, data were gathered in Namibia from two differently-sized congeneric species of orthopteran (La Greca & Messina 1989), the pest *Acanthoplus discoidalis* (Walker) and *Acanthoplus longipes* (Charpentier).

*Acanthoplus discoidalis* occurs in areas with low but thick vegetation. It is often found in the cultivated parts North-Eastern Namibia, where it is considered a significant pest, feeding on cereals such as sorghum and millet. However, this species is also opportunistically carnivorous (La Greca & Messina 1989), and has been recorded preying on other insects and even bird nestlings, such as the Red-billed Quelea (Cheke *et al*. 2003).

*Acanthoplus longipes* prefers semi-arid and arid habitats: its distribution includes an eastern strip that follows the course of the coastline and is widening from the North to the South (Cigliano *et al*. 2020). Large groups of individuals are often found on acacia shrubs, on which they actively feed grazing and sucking their leaves. They are primarily phytophagous albeit occasionally omnivorous (La Greca & Messina 1989). Also this species may become carnivorous, mostly of injured or dead conspecific individuals.

Our aim was to test the validity of statistical assumptions underlying the animal trait distributions, in order to verify which traits are statistically meaningful and verify the assumptions of the underlying hypotheses: are the trait distributions of body measurements primarily significantly influenced by sex (intraspecific variance) or by species (interspecific variance)? Our next questions were: can we identify any other major environmental characteristics that influences the morphological trait distribution, for example, vegetation canopy type? In addition, to what extent do shifts in the trait distribution influence the species from a behavioural perspective (locomotory implications, mating, etc.)? We therefore undertook an analysis of traits of both *Acanthoplus* species as a function of sex, species and habitat preference.

## METHODS

During March 2012, we collected 60 adults of *Acanthoplus longipes* (30 males and 30 females) near Keetmanshoop (25°52’1.2”S 18°06’12.5”E) and 46 adults of *Acanthoplus discoidalis* (20 males and 26 females) near Otjiwarongo (20°25’50.6”S 16°40’10.8”E). For each specimen we measured five functional traits using a Borletti caliper (measurement error of 0.02 mm). Specifically (see Fig. 1), we measured the total Body Length (henceforth BL) from the tip of the head to the end of the abdomen (excluding the ovipositor), the maximal pronotal width (PRW) as proxy for the entire body width, the maximal pronotal length (PRL), and finally the length of femur and tibia of the third pair of legs (FE_3_ and TI_3_, respectively) for both *A. discoidalis* (D) and *A. longipes* (L).

**Fig. 1.**
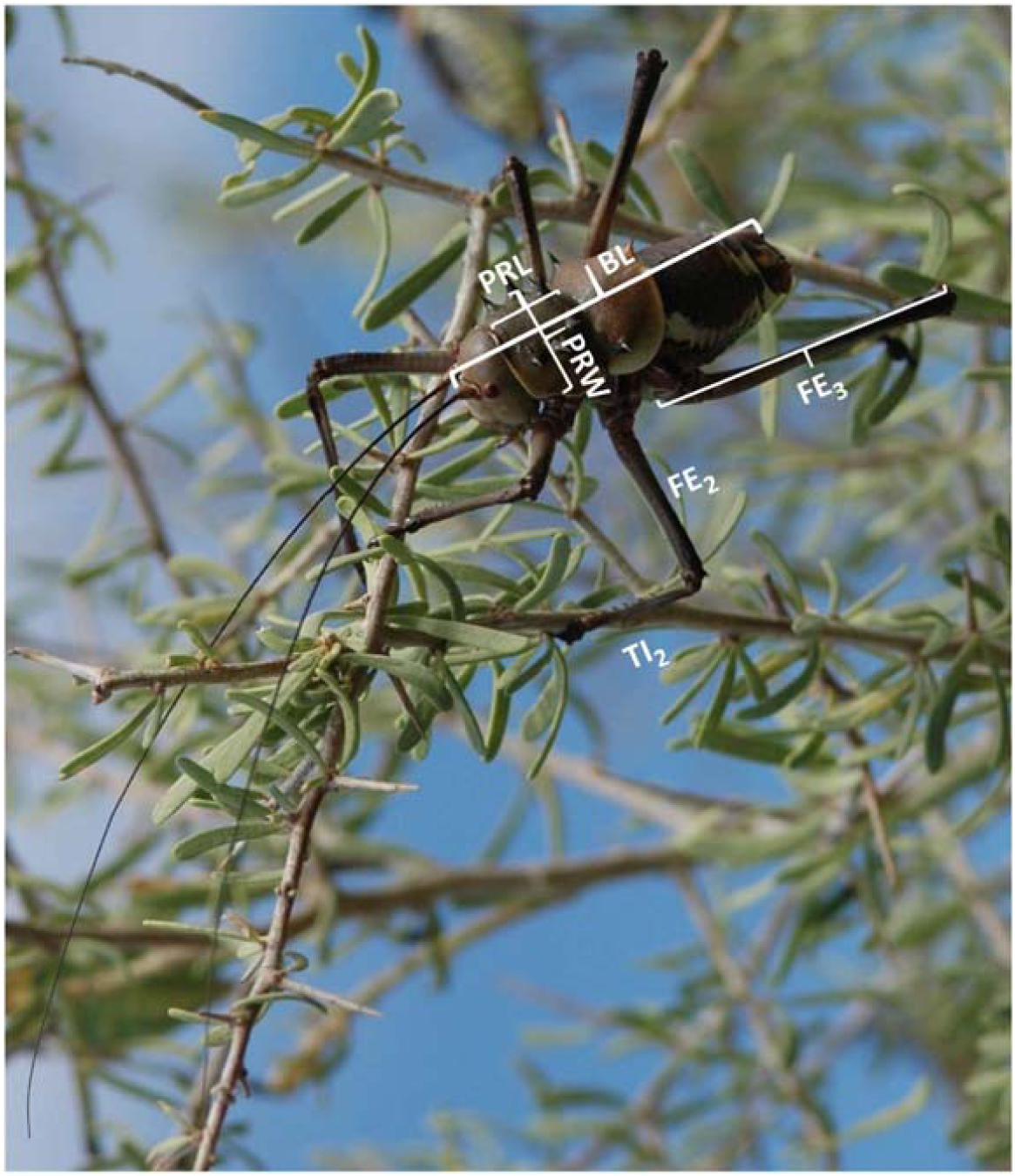
The field-measured functional traits for *Acanthoplus* body (BL, PRW) and hind legs (FE_3_, TI_3_ not shown) but see the second pair, FE_2_ and TI_2_, for comparison.

We estimated the body mass (BM) of our specimens using the most comprehensive model (the so-called “Model LWTR for the Tropics”) by Sohlström *et al*. (2018) which considers body Length, Width, Taxonomic group and geographic Region according to the following equation: log_10_(BM) = a_taxon_ Region + b_length_ Taxon Region × log_10_(Length) + b_width_ Taxon Region × log_10_(Width) where a_taxon_ = −0.117, b_lenght_ = 1.001, and b_width_ = 1.673 (Sohlström *et al*. 2018).

The Body Volume (BV) for *Acanthoplus* was also quantified by approximating the insect shape to a box constrained by the length and the width that can control BV according to the following equation: BV = BL × PRW^2^ (in mm^3^). If PRW data were unavailable, as in the case of Bidau & Martínez (2018) online data, we used own conversion factors to change BM into BV according to our own measurements: BV = 81.098 × BM −1014.879 (Bidau & Martínez 2018).

To evaluate if a species-specific correlation exists for the hind legs inside the genus *Acanthoplus*, we performed a regression relationship between femur and tibia and the total length of the third pair of legs calculated as the sum of femur and tibia, FE_3_+TI_3_. To test for inter- and intraspecific differences we used a one-way ANOVA using body length, length and width of pronotum and femur and tibia lengths as empirical variables and body mass estimates as derived variable (α = 0.01). Post-hoc comparisons were conducted using Dunnett’s T3 test. In order to evaluate the existence of any interspecific separation, we performed a Discriminant Function Analysis (DFA) considering the aforementioned functional traits as empirical variables. These statistical analyses were all performed using SPSS (vers. 21 for Windows).

Finally, we visualised the trait distribution using violin plots as realised with the “ggplot2” program by the “geom_violin()” utility in R-3.5.1.

## RESULTS

The comparison of males (m) and females (f) between the two species demonstrated a significant signal for five of the six traits (Table 1, Fig. 2). This indicates that, apart from body length (*F* = 0.488, *P* = 0.488 for males and *F* = 0.533, *P* = 0.468 for females), other measured traits were significantly different between D and L (Fig. 2). In particular, the pronotal traits, both Dm vs. Lm and Df vs. Lf were highly significantly different (*P* < 0.001). Tibia and femur lengths (TI_3_ and FE_3_, respectively) appear to be species-specific and are almost non-overlapping between *A. discoidalis* and *A. longipes* (Fig. 2, bottom). The correlation and regression analyses between TI_3_ and FE_3_ highlight strong relationships inside the genus since TI_3_ as sole predictor contributes for 60% to the expected total length of the hind legs.

**Table 1.** Empirical trait values for body length (BL), pronotal width (PRW), pronotal length (PRL), tibia length (TI_3_) and femur length (FE_3_) of our groups: the pest *A. discoidalis* males (Dm) and *A. discoidalis* females (Df), and the congeneric *A. longipes* males (Lm) and *A. longipes* females (Lf). Data are expressed as mean ± standard deviation in the first line; followed in the second line by the range values from 10^th^ and the 90^th^ percentage (shown in brackets).

|  | Dm | Df | Lm | Lf |
| --- | --- | --- | --- | --- |
| BL | 37.24 ± 4.21<br>(30.04 – 41.59) | 37.47 ± 5.07<br>(29.79 – 43.22) | 36.54 ± 2.86<br>(32.91 – 40.90) | 38.29 ± 3.14<br>(34.00 – 41.92) |
| PRW | 11.19 ± 0.48<br>(10.42 – 11.95) | 11.41 ± 0.59<br>(10.64 – 12.06) | 12.99 ± 0.58<br>(12.40 – 13.86) | 12.74 ± 0.67<br>(11.82 – 13.60) |
| PRL | 14.58 ± 0.79<br>(13.16 – 15.57) | 13.95 ± 0.84<br>(12.44 – 15.00) | 17.27 ± 0.99<br>(16.01 – 18.97) | 16.84 ± 1.80<br>(12.82 – 18.09) |
| TI<sub>3</sub> | 25.74 ± 1.40<br>(24.03 – 27.96) | 26.28 ± 1.60<br>(23.94 – 28.75) | 33.25 ± 1.68<br>(31.50 – 35.49) | 34.20 ± 1.58<br>(32.06 – 36.38) |
| FE<sub>3</sub> | 22.15 ± 1.29<br>(20.70 – 24.17) | 22.73 ± 1.04<br>(21.00 – 24.12) | 27.03 ± 1.31<br>(25.03 – 28.94) | 27.73 ± 1.31<br>(25.79 – 29.28) |

**Fig. 2.**
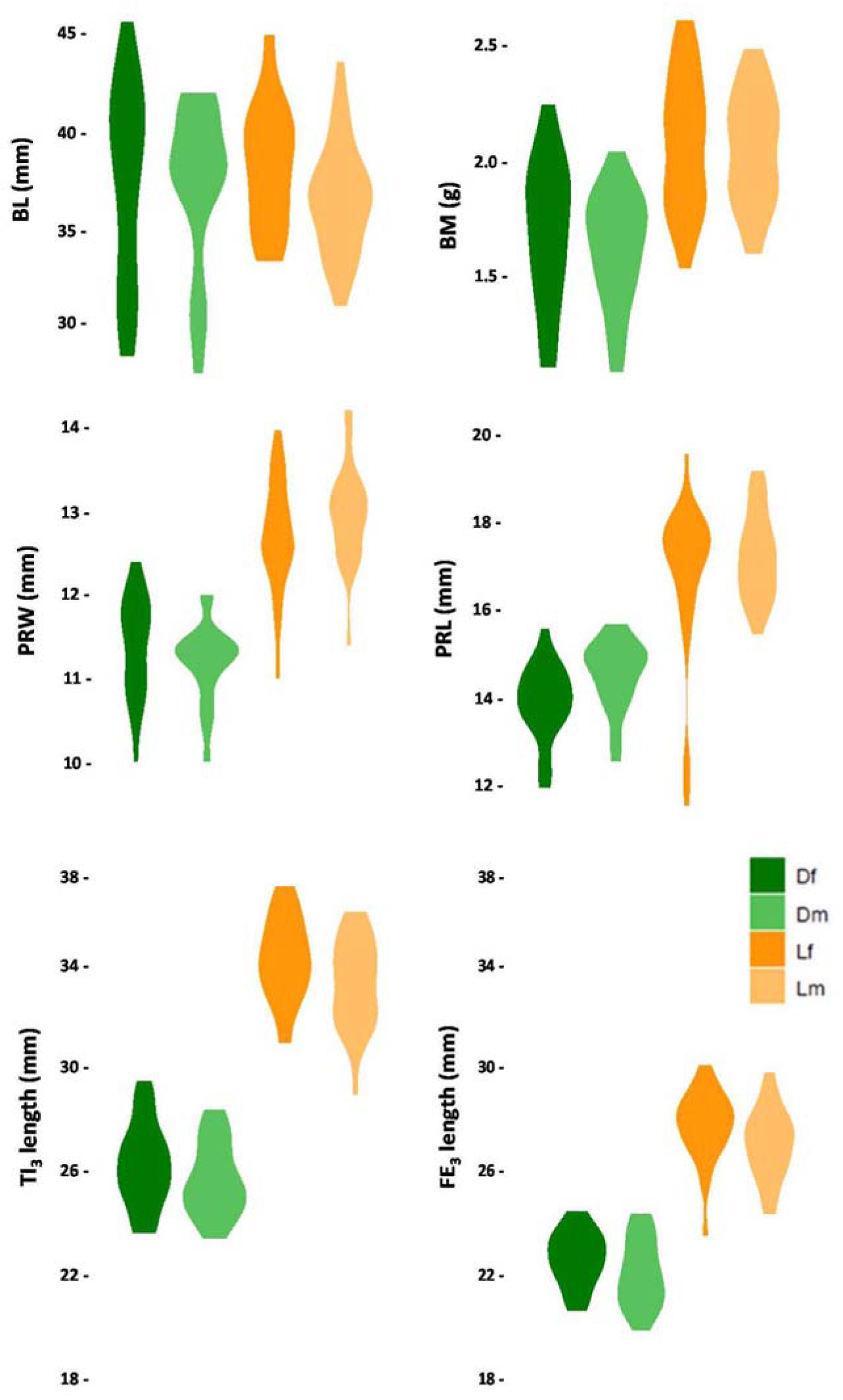
The trait distributions of body size and mass (upper panel), pronotal sizes (medium panel) and leg size (bottom panel) of our *Acanthoplus* specimens. The pest Dm = *A. discoidalis*, males, and Df = *A. discoidalis*, females, and the congeneric Lm = *A. longipes*, males, and Df = *A. longipes*, females. Trait abbreviations as in Table 1.

The first two discriminant functions (DFs) together account for 99% of the total variance and clearly separate the two species morphologically. The first DF has locomotory relevance as indicated by differences in tibia and femur lengths whilst the second DF reflects differences in size traits as captured by body and pronotal lengths (Table 2). It appears clear that the length traits of body and pronotum (DF2) and femur and tibia (DF1) are the relevant factors in differentiating between these two species, as body width was not significantly different (DF3, Wilks’ λ = 0.931, *P* = 0.066). In particular, all the centroids of our groups fall in four distinct quadrants (Fig. 3): the males of *A. longipes* share for both their legs and their size a DF>0 and fall into the first quadrant, and the females of the same species share DF>0 for the legs but DF<0 for the size and fall into the second quadrant. In contrast, the females of *A. discoidalis* share for both their legs and their size a DF<0 and fall into the third quadrant, and the males of the same species share DF<0 for the legs but DF>0 for the size and fall into the fourth quadrant. Our analyses unequivocally separate the pest species *A. discoidalis* from its congeneric species.

**Table 2.**
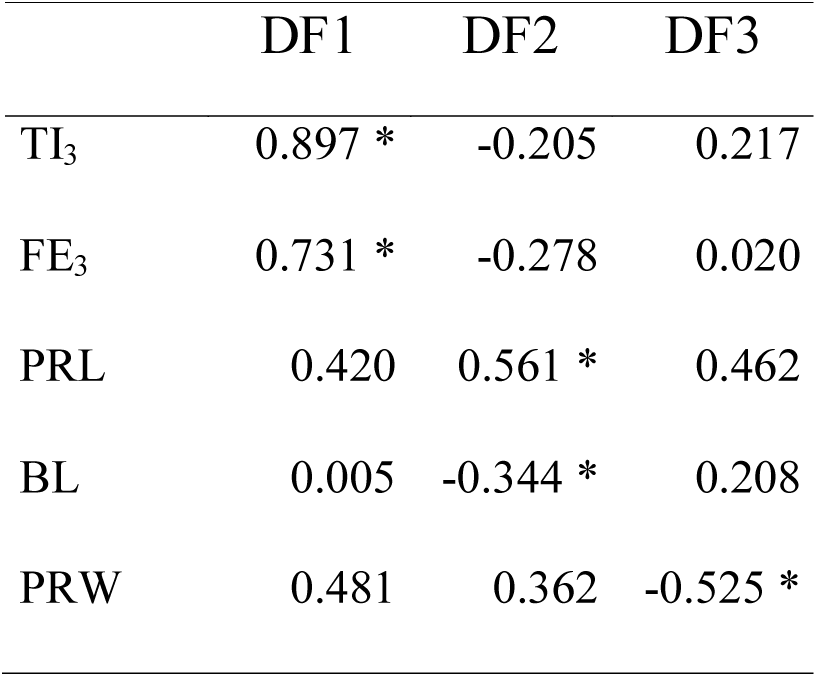
Pooled within-groups correlations between discriminating traits and standardized canonical discriminant functions (DFs); the traits are ordered by absolute size of correlation within DFs. Asterisks represent the largest absolute correlation between each trait and any DF. Trait abbreviations as in Table 1.

**Fig. 3.**
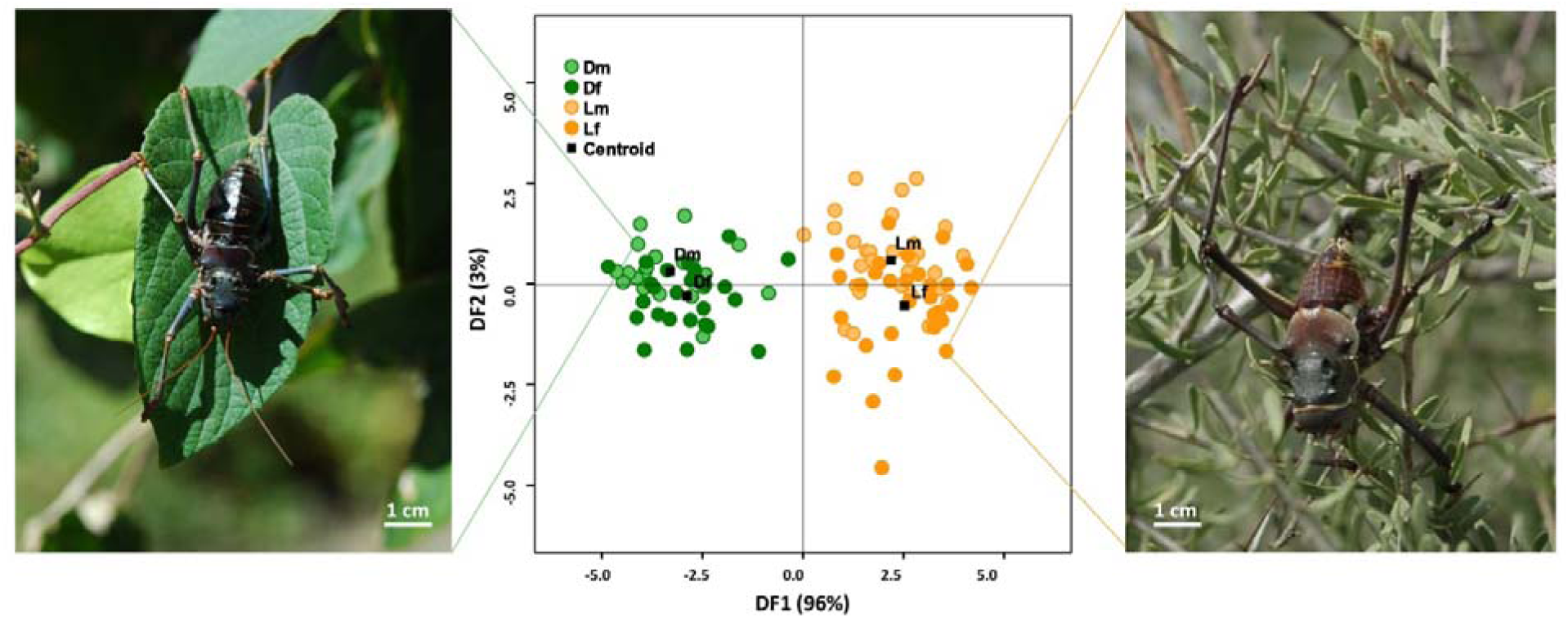
Scatter plot of discriminant function analysis (DFA) based on the first two components, taking into consideration femur and tibia (DF1) and length traits of body and pronotum (DF2). Colours and symbols as in the previous figure. Specimens of *A. discoidalis* (on the left) and *A. longipes* (on the right) are shown

The recently published data by Bidau & Martínez (2018) provided BL and FE_3_ trait values for 50 African tettigoniids (25 for each gender). A multiple comparison of body mass, body length and femur length between sexes resulted in highly significant results (females BM *F* = 23.20, BL *F* = 117.17 and FE_3_ *F* = 31.65; males BM *F* = 36.84, BL *F* = 121.73 and FE_3_ *F* = 47.13, with *P* < 0.0001 in all the six cases), pointing to intraspecific variation. Hence we plotted the FE_3_ length as a function of estimated insect volume of our genus in comparison to other tettigoniids (Fig. 4).

**Fig. 4.**
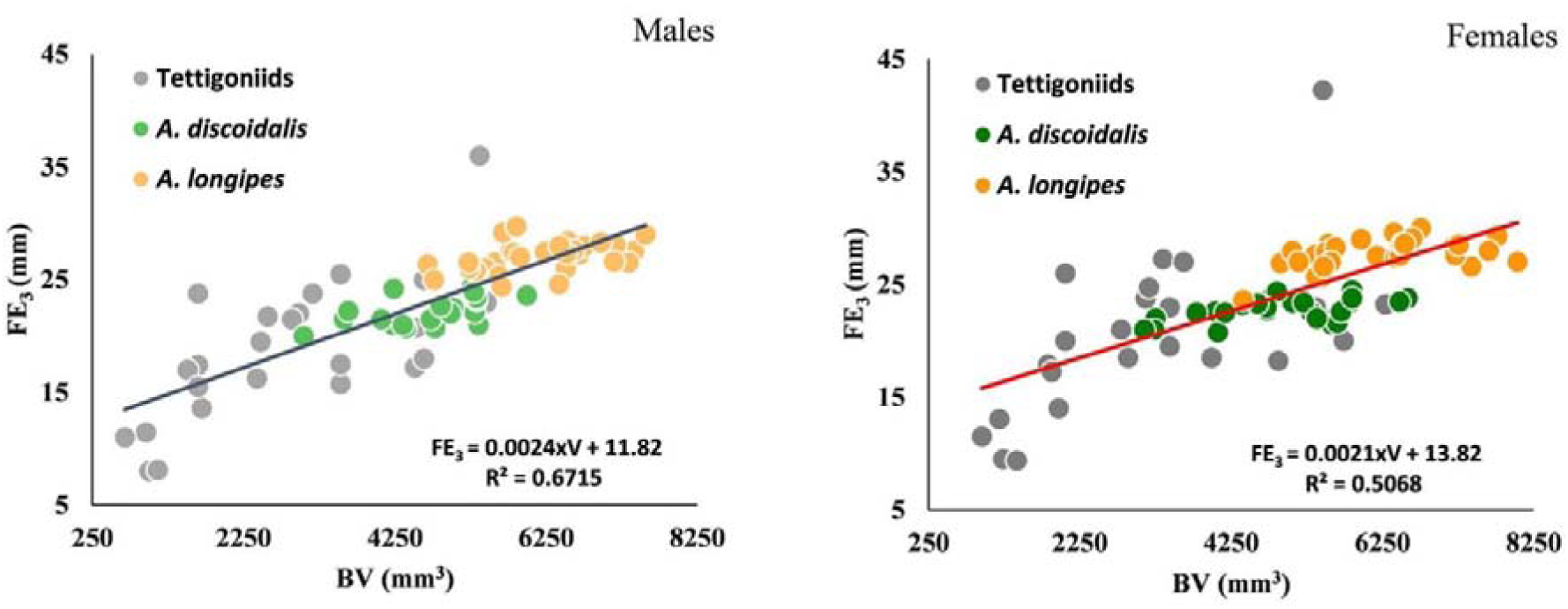
Allometric scaling of femur length (FE_3_) as linear function of body volume (BV) in males (left panel) and females (right panel) of Namibian *Acanthoplus* specimens (same colours as in the previous figures) in comparison to African tettigoniids (grey-filled circles) as derived from the online dataset by Bidau & Martínez (2018).

## DISCUSSION

Behavioural differences between species can provide more information than can single physiological and/or morphological aspects (Keenleyside 1979; Bateman & Fleming 2008). As behaviour represents the adaptive response of individual of a species to changing environmental features, one can assume that any change in a behavioural pattern may be reflected in a stronger selection for a corresponding morphological change (Gwynne 1981; Peters 1983; Pough *et al*. 1992; Bertossa 2011). Nevertheless, variation in behavioural patterns do not follow a strong allometric or isometric relationship as morphological structures do (cf. Green *et al*. 2001). The necessity to find a link between behavioural changes and morphological traits has become more and more urgent in evolutionary divergence and coevolution given the high data amount that became available for modelling (Kalinkat *et al*. 2015; Willmer *et al*. 2009). Examples come out by vocalization patterns and escape mechanisms (Dial *et al*. 2008 and Bateman & Fleming 2008, respectively).

In most ectotherms, females tend to be longer than males (Fairbairn 1997; Blanckenhorn *et al*. 2007). Rensch’s rule (1950, 1959), that any sexual size dimorphism increases with BL in the species where the males are larger than their mates and decreases with BL in the opposite case, was demonstrated in our specimens. In particular, we found no statistical evidence for sexual size dimorphism for BL but only for BM (Fig. 2). As the latter trait is a function of body length and pronotal width, we think that, in comparison to BL, the pronotal width is the relevant trait in explaining the interspecific differences in the calculated BM values. On the other hand, pronotal dimensions are strictly related to the stridulatory organs in males and these two Namibian species of *Acanthoplus* perform quite distinct calling songs (Conti & Viglianisi 2005; Kowalski & Lakes-Harlan 2011). Evidence of the positive trait scaling of the wing area with body mass and body size have been shown for the bush cricket *Poecilimon ampliatus* (Anichini *et al*. 2017).

Foreseen breakthroughs lie in the development of novel, trait-driven frameworks for comprehensive ecological evaluation. Our study clearly indicates that the entire trait distribution is significantly more different at interspecific than at intraspecific levels and considers the implications of trait-driven behavioural patterns in two congeneric *Acanthoplus* species.

We argue that the larger-sized *A. longipes* might reflect in its locomotory traits a kind of adaptation that improves the possibility to find a mate grasping into the branches of the shrub they inhabit. In contrast, the shorter-legged pest *A. discoidalis* reflects in its locomotory traits a successful attempt to compensate for the high energy loss due to their longer-pulsed calling songs (Kowalski & Lakes-Harlan (2011), but see Bailey *et al*. (1993) for bush crickets) and the high energy input due to elevate temperatures of the soil surface (cf. Martin *et al*. 2000) with a slower movement across the cereal fields. This may be a matter of differences in philosophy, but we consider these to have strong implications at gender level. It might in fact seem to be hard to choose for either a sexual selection on males or a reproductive selection on females, but the pointed cerci of the males that allow a much firmer grip of the females (Irish 1992) suggest here a sexual selection on males, making their calling songs very relevant. Kowalski & Lakes-Harlan (2011) analysed the calling songs of these species across a temperature gradient without evidence of any temperature effect on their songs.

All our analyses not only unequivocally separate the pest species from its congeneric species, but also show that these *Acanthoplus* ground crickets are much larger than other African tettigoniids (Fig. 4). It is, furthermore, remarkable that tettigoniids have on average a nitrogen content of 11.2% in their tissues, much higher than the overall nitrogen content of 9.9% of other insects (Fagan *et al*. 2002). Being even larger, our *Acanthoplus* specimens demand from the foliar tissues they consume a much higher nutrient quality. Indeed, we argue that despite their omnivory, our tettigoniids have a remarkable difference in their food preference. The larger-sized *A. longipes* preferably feeds on *Acacia* leaves that as N_2_-fixers have a much higher than average nitrogen percentage whilst the pest *A. discoidalis* is mostly feeding on cereals with a much lower nitrogen percentage (respectively 3.5% vs. 2.1% of the foliar tissues of the host plants according to Mulder *et al*. 2013). This difference in nitrogen content allows our species to reach differences in mass that, in turn, needs different size in their locomotory traits-This adaptation is of highest relevance because both species cannot swarm.

Summarising, functional traits are known to be a valuable proxy in ecology and relevant traits are much more than body size alone. Still, different interspecific aspects of size contribute by far the most to the results of our discriminant analysis (Fig. 3), highlighting the relevance of functional traits as real evolutionary clumps. Notwithstanding the great potential of existing trait-based models, up to now, too many animal ecologists seem unaware of the benefits traits provide. These analyses convincingly demonstrate that the presumed relevance of gender-specific trait-distributions in literature is not always correct and call into question some of the invoked links between the individual-based traits and the response of the population structures.

## Authors’ contributions

E.C. and G.C. conducted the fieldwork (permit #1675/2012); E.C. and C.M. conceptualised the study and contributed to data curation; E.C. contributed to the modelling and software; E.C., G.C. and C.M. contributed to the writing of the manuscript.

## Acknowledgements

This research was partially funded by the Ministry of Education, Universities and Research, Rome, Italy. We thank Alessandro Marletta and the Ministry of Environment and Tourism, Windhoek, Namibia, and are grateful to the Editor and one anonymous referee for their comments.

